# A conserved motif in Pch2 regulates its localization and meiotic function in *Saccharomyces cerevisiae*

**DOI:** 10.64898/2026.06.04.730238

**Authors:** Esther Herruzo, Sara Téllez, Beatriz Santos, Pedro A. San-Segundo

## Abstract

The *Saccharomyces cerevisiae* Pch2 protein is a conserved meiotic AAA+ ATPase whose activity must be tightly regulated to ensure proper chromosome dynamics during meiotic prophase I. Its function relies on remodeling the HORMA-domain protein Hop1, promoting conformational transitions that are essential for chromosome axis organization, checkpoint signaling, and recombination control. Here, we identify threonine 428 (T428), located within a conserved threonine-glutamine (TQ) putative phosphorylation motif, as a critical regulatory residue of Pch2. We found that, in *zip1Δ* cells, the meiotic recombination checkpoint response is partially or completely abolished in the *pch2-T428A* and *pch2-T428D* mutants, respectively. Both mutations alter Pch2 subcellular localization, leading to its increased nuclear accumulation; however, forced nuclear exclusion of Pch2-T428A, but not Pch2-T428D, restores the *zip1Δ* meiotic block, indicating an additional effect of the T428D substitution on checkpoint function beyond subcellular distribution. Analysis in synapsis-proficient strains reveals that this residue also plays a critical role in coordinating Hop1 chromosomal enrichment with Mek1 activation along the synaptonemal complex. In contrast to *pch2Δ* or the ATPase-defective *pch2-E399Q* mutant, introduction of a negative charge at the 428 position uncouples Hop1 accumulation from its phosphorylation, preventing Mek1 activation despite robust Hop1 association with meiotic chromosomes. These findings support emerging models in which Pch2 regulates Hop1 to control not only its chromosomal abundance, but also the maintenance of sufficient levels of Hop1 in a phosphorylation-competent conformation, thereby ensuring proper checkpoint signaling and faithful meiotic progression.

## INTRODUCTION

Meiosis is a specialized type of cell division essential for sexually reproducing organisms. It reduces chromosomal number by half, enabling fertilization and new allele combinations. Following DNA replication, during a tightly regulated prophase I, homologous chromosomes pair, synapse and recombine, to finally segregate towards opposite poles in the first meiotic division. In the second division, sister chromatids separate, producing four haploid gametes (Borner et al. 2023; Zickler and Kleckner 2023). Meiotic recombination initiates with the introduction of DNA double strand breaks (DSBs) in the replicated DNA by the topoisomerase-like protein Spo11 (Arter and Keeney 2024). These breaks are processed to generate single-stranded DNA (ssDNA) that is proficient to incorporate the recombinases Rad51 and Dmc1, which promote homology search and pairing of homologous chromosomes (Brown and Bishop 2014). To stabilize these interactions, a proteinaceous structure known as the synaptonemal complex (SC) is assembled between homologs (Lascarez-Lagunas et al. 2020; Cesar and Kim 2026). In *Saccharomyces cerevisiae*, the SC consists of two axial cores, or lateral elements (LEs), formed by Red1, Hop1 and Rec8-containing cohesin, and a central region composed of Zip1 transverse filaments together with the Ecm11 and Gmc2 proteins that constitute the central element. The Rec8 cohesin complex binds and extrudes DNA forming chromatin loops (Klein et al. 1999; Schalbetter et al. 2019). Red1 anchors to the base of these loops through the assembly of coil-coiled filaments and directly recruits Hop1 to the axis core. This chromosomal organization is crucial for meiotic recombination because it provides the structural framework that accommodates DSB formation, signaling, and repair (Hollingsworth et al. 1990; Sym et al. 1993; Smith and Roeder 1997; Sun et al. 2015; Heldrich et al. 2022; Prince and Martinez-Perez 2022). A subset of meiotic DSBs is repaired as crossovers (COs), leading to genetic exchange and the establishment of physical connections between homologs, called chiasmata, which ensure correct orientation and segregation of chromosomes at meiosis I (San-Segundo and Clemente-Blanco 2020; Raghavan and Hochwagen 2025).

To minimize genome instability and reduce the risk of aneuploidy arising from defects in these processes, meiotic cells employ a conserved surveillance system known as the meiotic recombination checkpoint. This pathway monitors the progress of recombination and synapsis and imposes a prophase I arrest or delay when unrepaired recombination intermediates persist (Subramanian and Hochwagen 2014). In *zip1Δ* mutants, where synapsis fails and unrepaired DSBs accumulate, the Mec1 kinase (ortholog of human ATR) is recruited to resected DSB ends via its partner Ddc2 (Refolio et al. 2011; Sampathkumar et al. 2024). Mec1 phosphorylates Hop1 at multiple SQ/TQ motifs; among these, phosphorylation of Hop1-T318 is critical for the subsequent recruitment and activation of the meiosis-specific effector kinase Mek1 (Carballo et al. 2008; Penedos et al. 2015). Activated Mek1 then phosphorylates the Ndt80 transcription factor, inhibiting its activity and preventing the expression of key meiotic cell-cycle regulators, such as *CLB1* and *CDC5* (Chen et al. 2018). Together with Swe1-dependent inhibitory phosphorylation of Cdc28/Cdk1, this signaling network establishes the meiotic prophase I arrest (Leu and Roeder 1999; Gonzalez-Arranz et al. 2018; Gonzalez-Arranz et al. 2026). Beyond cell-cycle control, Mek1 also modulates the recombination outcome by phosphorylating Rad54 and Hed1, thereby imposing an interhomolog bias that prioritizes DSB repair using the homologous chromosome rather than the sister chromatid (Niu et al. 2009; Callender et al. 2016).

The critical role of the AAA+ ATPase Pch2 in this checkpoint signaling cascade is to remodel Hop1 conformation, a step required to maintain adequate levels of Red1-dependent Hop1-T318 phosphorylation and proper Hop1 localization to chromosome axes (Lo et al. 2014; Herruzo et al. 2016; Herruzo et al. 2021). Pch2 (TRIP13 in mammals) was originally identified in *S. cerevisiae* as a pachytene checkpoint factor that enforces meiotic arrest in response to synapsis defects (San-Segundo and Roeder 1999). However, it is currently known that Pch2 regulates a wide variety of molecular functions during meiosis across different organisms (Bhalla 2023). Pch2 assembles into hexameric rings (Chen et al. 2014) and contains a conserved catalytic domain with ATP binding and hydrolysis motifs, as well as a regulatory N-terminal domain containing nuclear localization (NLS) and nuclear export (NES) signals (Herruzo et al. 2019; Herruzo et al. 2021; Herruzo et al. 2023). In most systems, the hexameric Pch2 utilizes the energy derived from ATP hydrolysis to mechanically remodel meiotic HORMAD proteins (for Hop1, REV7, and MAD2) from a “closed” to an “open” conformation to control their function, thereby coordinating meiotic homolog pairing, synapsis and recombination (Rosenberg and Corbett 2015; Vader 2015; Ye et al. 2017; West et al. 2018; Puchades et al. 2020; Gu et al. 2022; Prince and Martinez-Perez 2022; Russo et al. 2023). In contrast to other organisms (Eytan et al. 2014; Balboni et al. 2020; Giacopazzi et al. 2020), in *S. cerevisiae* no cofactor appears to be required for the remodeling action of Pch2 on Hop1.

In addition to the critical function of Pch2 in the *zip1*Δ checkpoint, it also performs important functions during unperturbed meiosis. Recruitment of Pch2 to synapsed chromosomes primarily depends on Zip1, but other factors such as RNAPII-dependent transcription, Top2, Nup2, and chromatin modifications driven by Sir2 and Dot1, also influence its chromosomal distribution (San-Segundo and Roeder 1999, 2000; Ontoso et al. 2013; Cavero et al. 2016; Subramanian et al. 2019; Cardoso da Silva et al. 2020; Heldrich et al. 2020). Recruitment of Pch2 to the SC as synapsis proceeds drives the release of Hop1 from the axes (San-Segundo and Roeder 1999; Borner et al. 2008; Herruzo et al. 2016), with several functional consequences. Because Hop1 promotes DSB formation, its removal contributes to shutting down the DSB-forming machinery, thereby limiting further DSB formation once homologs are synapsed (Thacker et al. 2014; Mu et al. 2020). Consistently, Hop1 exists in two different states: a chromatin-bound DSB-promoting form that is relatively resistant to Pch2, and a DSB-inactive state that is more accessible to Pch2-dependent removal (Milano et al. 2024). However, Pch2-dependent Hop1 depletion is not uniform, as Hop1 is retained at telomere-proximal end-adjacent regions (EARs), which maintain recombination competence despite synapsis (Subramanian et al. 2019). In addition, Hop1 release leads to local loss of Mec1-dependent Hop1/Mek1 signaling, relieving interhomolog bias and enabling timely repair of meiotic DSBs (Subramanian et al. 2016). Thus, Pch2 couples synapsis to the spatiotemporal coordination of recombination and meiotic progression, ensuring that recombination is established under interhomolog bias and subsequently completed once homolog engagement has occurred.

Pch2 also regulates recombination in specialized chromosomal regions such as the heterochromatin-like ribosomal DNA (rDNA) array. Pch2 is highly enriched in the nucleolus, where it excludes Hop1 from the rDNA, thereby suppressing recombination within this highly repetitive region and maintaining rDNA stability. This nucleolar function of Pch2 is mechanistically distinct from its role at euchromatic chromosome regions and is not required for meiotic checkpoint signaling. Consistent with this, Pch2 recruitment to the rDNA depends on its interaction with the origin recognition complex (ORC) component Orc1, which prevents Hop1 accumulation and unwanted recombination in the nucleolus but is dispensable for checkpoint function (San-Segundo and Roeder 1999; Vader et al. 2011; Herruzo et al. 2019; Villar-Fernández et al. 2020).

In addition to its chromosomal and nucleolar roles, a cytoplasmic pool of Pch2 has been identified and is essential for meiotic checkpoint activation. Genetic and cytological evidence indicate that this extranuclear population promotes the generation of a signaling-competent pool of Hop1 that can be incorporated into chromosome axes and subsequently phosphorylated at T318, although the underlying mechanism remains unclear (Raina and Vader 2020; Herruzo et al. 2021). How cytoplasmic Pch2 influences Hop1 dynamics at a distance from chromosomes has not yet been resolved, but analogous regulatory features have been described in plants, where cytoplasmic PCH2 controls the availability and chromosomal loading of ASY1 (Hop1 ortholog) through remodeling-dependent mechanisms (Yang et al. 2020). Importantly, perturbations that alter the balance between nuclear and cytoplasmic Pch2 pools impair Hop1 chromosomal distribution and checkpoint signaling (Herruzo et al. 2021; Herruzo et al. 2023). Together, these observations indicate that Pch2 operates across multiple cellular and chromosomal compartments, and that a precise balance in its subcellular distribution is critical for its diverse meiotic functions.

To better understand this complex regulatory landscape, we analyzed the functional contribution of a conserved threonine-glutamine (TQ) motif within the Pch2 AAA+ domain, a potential target for phosphoregulation. Using a combination of mutational analysis, live-cell imaging, and chromosome cytology, we identify threonine 428 (T428) as a critical regulatory node. We find that introduction of a negative charge at this residue interferes with Pch2 nuclear export and uncouples its subcellular distribution from its catalytic function. As a consequence, Pch2-T428D fails to support robust meiotic checkpoint signaling and disrupts the coordination between Hop1 loading and Mek1 activation. By characterizing this residue, we provide new mechanistic insights into how the spatial distribution and functional activity of Pch2 ensure the fidelity of meiotic chromosome dynamics.

## MATERIALS AND METHODS

### Yeast strains and plasmids

The genotypes of yeast strains are listed in Table S1. All strains are in the BR1919 background (Rockmill and Roeder 1990). The *zip1Δ::LEU2, zip1Δ::LYS2*, *ndt80Δ::LEU2 ndt80Δ::kanMX3*, *pch2Δ::TRP1,* and *rad24Δ::TRP1* gene deletions were previously described (Sym et al. 1993; Sym and Roeder 1994; Lydall and Weinert 1997; San-Segundo and Roeder 1999; Tung et al. 2000; Herruzo et al. 2016). The *sml1Δ::kanMX6, mec1Δ::KIURA3, tel1Δ::hphMX6* and *hop1Δ::hphMX6* deletions were made using a PCR-based approach (Longtine et al. 1998; Goldstein and McCusker 1999). The *3HA-PCH2*, *P_HOP1_-GFP-PCH2, P_HOP1_-GFP-NES-PCH2, PIL1-GBP-mCherry::hphMX6* and *spo11-3HA-6His*::*kanMX4* constructs have been previously described (San-Segundo and Roeder 1999; Herruzo et al. 2021). The *pch2-K320A* and *pch2-E399Q* mutations were described in (Herruzo et al. 2016). The *pch2-T428A* and *pch2-T428D* mutations were introduced at the *PCH2* genomic locus using the *delitto perfetto* technique (Stuckey et al. 2011); they generate an *Hha*I and a *Bcl*I site, respectively, utilized for genotyping purposes during genetic crosses. The *HOP1-GFP* construct was amplified from an SK1 strain kindly provided by Neil Hunter (UC Davis) and inserted at the *HOP1* locus in BR strains. *ZIP1* was overexpressed using pSS343 plasmid previously described in (Sym and Roeder 1995).

All constructions and mutations were verified by PCR analysis and/or sequencing. The sequences of all primers used in strain construction are available upon request. All strains were made by direct transformation of haploid parents or by genetic crosses always in an isogenic background. Diploids were made by mating the corresponding haploid parents and isolation of zygotes by micromanipulation.

### Meiotic cultures and meiotic time courses

To induce meiosis and sporulation, BR strains were grown in 3.5 ml of synthetic complete medium (2% glucose, 0.7% yeast nitrogen base without amino acids, 0.05% adenine, and complete supplement mixture from Formedium at twice the particular concentration indicated by the manufacturer) for 20–24 h, then transferred to 2.5 ml of YPDA (1% yeast extract, 2% peptone, 2% glucose, and 0.02% adenine) and incubated to saturation for an additional 8 h. Cells were harvested, washed with 2% potassium acetate (KAc), resuspended into 2% KAc (10 ml), and incubated at 30°C with vigorous shaking to induce meiosis. Both YPDA and 2% KAc were supplemented with 20 mM adenine and 10 mM uracil. The culture volumes were scaled up when needed.

To score meiotic nuclear divisions, samples from meiotic cultures were taken at different time points, fixed in 70% ethanol, washed in phosphate-buffered saline (PBS) and stained with 1 μg/μl 4′,6-diamidino-2-phenylindole (DAPI) for 15 min. At least 300 cells were counted at each time point. Meiotic time courses were repeated several times; averages and error bars from at least three replicates are shown.

### Western blotting

Total cell extracts for Western blot analysis were prepared by trichloroacetic acid (TCA) precipitation from 3-ml aliquots of sporulation cultures, as previously described (Gonzalez-Arranz et al. 2026). Proteins were resolved by SDS-PAGE and then transferred to PVDF membranes. To resolve the phosphorylated forms of 3HA-Pch2 or Mek1, 10% SDS-PAGE gels with a 29∶1 ratio of acrylamide∶bisacrylamide containing 50 µM and 30 µM of the Phos-tag reagent (Wako Chemicals), respectively, and 75 µM MnCl_2_ were used. Gels were run on ice at 150 volts in a MiniProtean (Bio-Rad) for 5 h for 3HA-Pch2 and 100 volts for 4 h for Mek1. After running, gels were washed with 1 mM EDTA before transfer to PVDF membranes. Regular 8%, 10% or 12% gels (37.5:1 acrylamide∶bisacrylamide) were used for routine detection of 3HA-Pch2, Hop1-T318ph, Hop1, Hop1-GFP, H3-T11ph, H3 and Pgk1. The antibodies used are listed in Table S2. Antibodies were diluted in TBS 0.1% Tween containing 5% milk (for GFP, HA, Hop1, Mek1, Pch2 and Pgk1), 0.1% BSA (Bovine Serum Albumin) (for Hop1-T318ph), or 5% BSA (for H3 and H3-T11ph). PVDF membranes were incubated at 4°C overnight (primary antibodies), or at room temperature for 45 minutes (secondary antibodies). The Pierce ECL or ECL Plus reagents (ThermoFisher Scientific) were used for detection. Signals for Figures 1c, 1d, and 2c were captured on film (Amersham Hyperfilm ECL; GE Healthcare), whereas signals for Figures 3e, 5f, and 5h were acquired with a Fusion FX6 system (Vilber) and quantified using Evolution-Capt software (Vilber).

**Figure 1.**
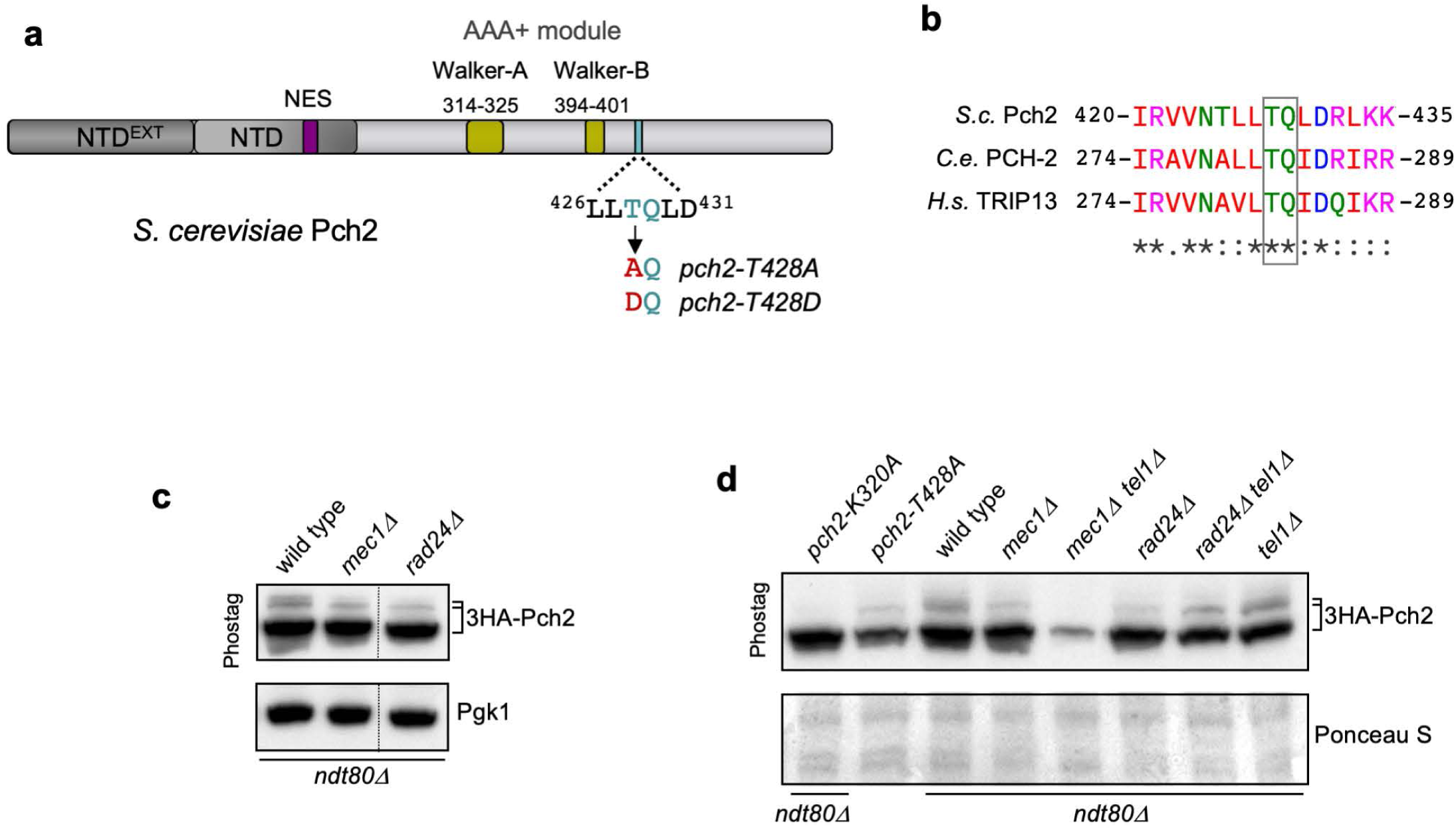
Pch2 contains a putative Mec1 phosphorylation motif. **a)** Schematic representation of the *S*. *cerevisiae* Pch2 protein (ScPch2) indicating the characteristic AAA+ ATPase motifs, the conserved N-terminal domain (NTD), the extended N-terminal domain exclusive of yeast Pch2 (NTD^EXT^) and the nuclear export signal (NES). The position and sequence of the conserved Mec1/Tel1 consensus phosphorylation site (TQ site) analyzed in this work is depicted (light blue) along with the modifications introduced (red) in the mutants generated in this work (*pch2-T428A* and *pch2-T428D*). **b)** ClustalW alignment of a portion of the protein sequences of Pch2 orthologs from *S. cerevisiae* (*S.c.* Pch2), *C. elegans* (*C.e*. PCH-2) and human (*H.s.* TRIP13) encompassing the conserved TQ motif (boxed). **(c-d)** Phos-tag western blot analyses of cell extracts from *ndt80*Δ-arrested cells after 24 h in meiosis, except the *pch2-T428A* lysate, which is from *NDT80* cells at 15 h in meiosis. Slow migrating putative phosphorylation forms of 3HA-Pch2 (detected with anti-HA antibodies) are marked on top of the basal band. Pgk1 or Ponceau S staining were used as loading controls in (c) and (d), respectively. Strains are: DP1193 (*3HA-pch2-K320A ndt80Δ*), DP1256 (*3HA-pch2-T428A*), DP1191 (wild type; *3HA-PCH2 ndt80Δ*), DP1205 (*mec1Δ sml1Δ 3HA-PCH2 ndt80Δ),* DP1184 (*mec1Δ sml1Δ tel1Δ 3HA-PCH2 ndt80Δ*), DP1197 (*rad24Δ 3HA-PCH2 ndt80Δ*), DP1183 (*rad24Δ tel1Δ 3HA-PCH2 ndt80Δ*) and DP1252 (*tel1Δ 3HA-PCH2 ndt80Δ*).

**Figure 2.**
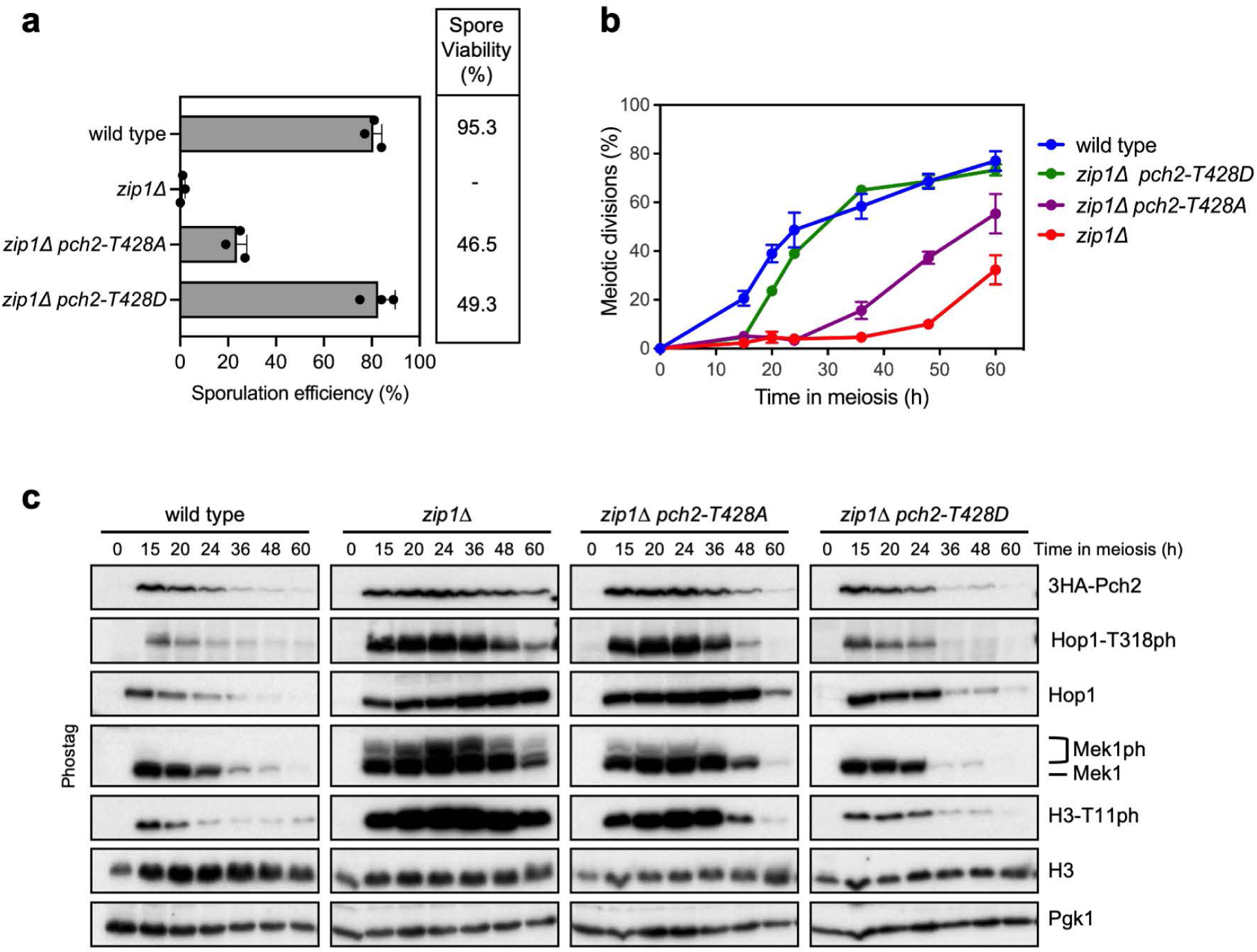
The meiotic recombination checkpoint is strongly impaired in the *zip1*Δ *pch2-T428D* mutant. **a)** Sporulation efficiency, assessed by microscopic counting of asci, was examined after 3 days on sporulation plates. Error bars, SD; n = 3. At least 300 cells were counted for each strain. Spore viability, assessed by tetrad dissection, is also shown. At least 36 tetrads were dissected for each strain. **b)** Time-course analysis of meiotic nuclear divisions; the percentage of cells containing two or more nuclei is represented. Error bars: SD; n=3. At least 300 cells were scored for each strain at every time point. **c)** Western blot analysis of Pch2 (detected with anti-HA antibodies), Hop1 and histone H3 production, and checkpoint signaling (Hop1-T318ph, Mek1ph and H3-T11ph) during meiotic time courses of the indicated strains. For Mek1ph, a Phos-tag gel was used. Pgk1 was used as a loading control. Strains are: DP1151 (wild type; *3HA-PCH2*), DP1152 (*zip1Δ 3HA-PCH2*), DP1257 (*zip1Δ 3HA-pch2-T428A*) and DP1333 (*zip1Δ 3HA-pch2-T428D*).

**Figure 3.**
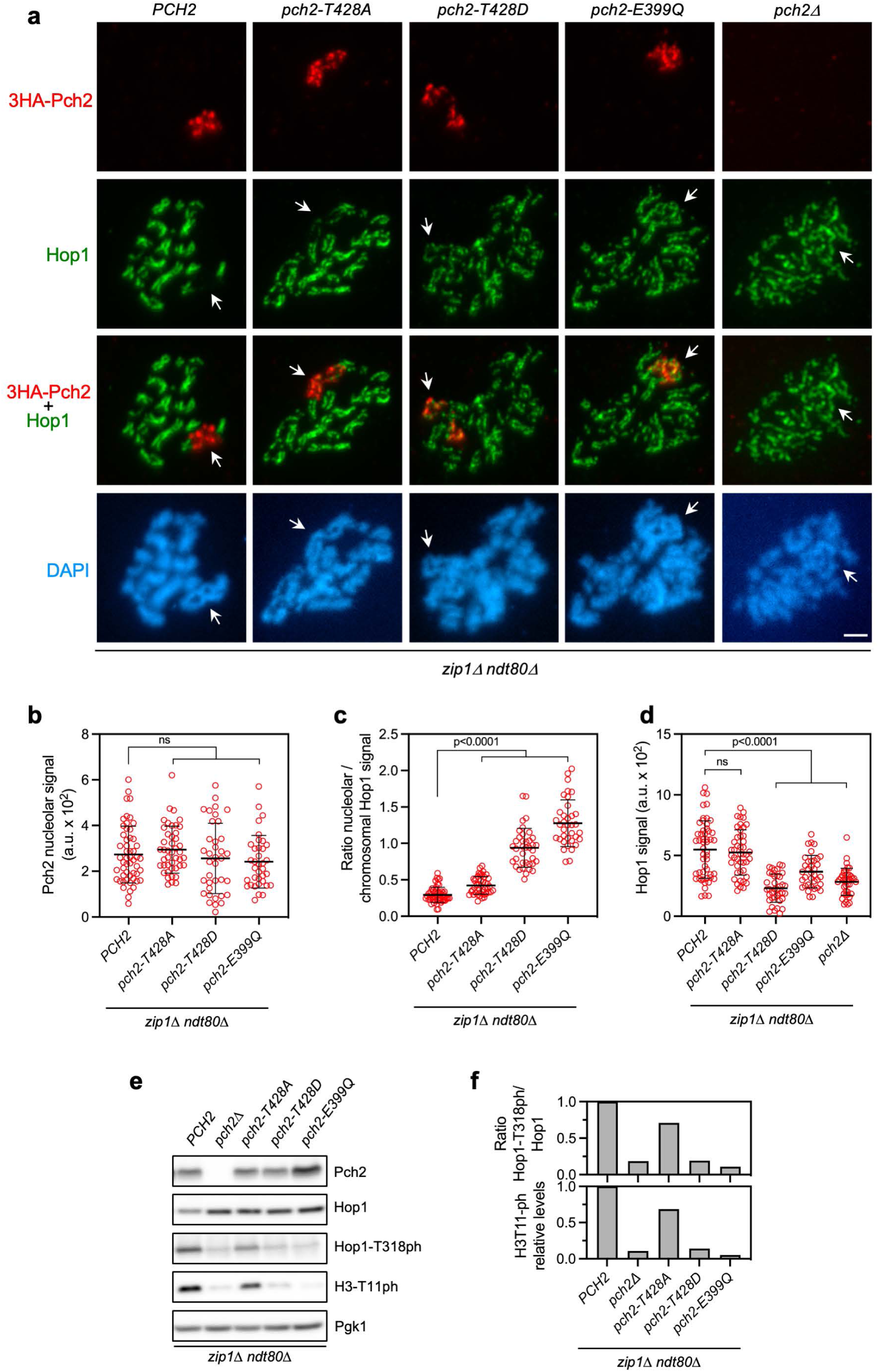
Hop1 localization and phosphorylation are defective in the *zip1*Δ *pch2-T428D* mutant. **a)** Immunofluorescence of spread meiotic chromosomes from prophase-I arrested *zip1*Δ *ndt80*Δ strains, stained with anti-HA (to detect 3HA-Pch2; red) and anti-Hop1 antibodies (green), and with DAPI (blue). Spreads were prepared 24 h after meiotic induction. Representative nuclei are shown. Arrows point to the rDNA region. Scale bar, 2 μm. **b-d)** Quantification of the nucleolar Pch2 signal (b), the ratio of nucleolar to chromosomal Hop1 signal (c), and the total Hop1 signal (d) for the experiment shown in (a). Error bars, SD; a.u., arbitrary units. **e)** Western blot analysis of Pch2 (detected with anti-HA antibodies), Hop1 and phosphorylation of Hop1-T318 and H3-T11. Pgk1 was used as a loading control. **f)** Quantification of the normalized ratio of Hop1-T318ph to Hop1, and of H3T11ph relative levels from the experiment in (e). Strains are: DP1190 (*zip1Δ ndt80Δ 3HA-PCH2*), DP881 (*zip1Δ ndt80Δ pch2Δ*), DP1285 (*zip1Δ ndt80Δ 3HA-pch2-T428A*), DP1400 (*zip1Δ ndt80Δ 3HA-pch2-T428D*), and DP1302 (*zip1Δ ndt80Δ 3HA-pch2-E399Q*).

### Cytology

Immunofluorescence of chromosome spreads was performed essentially as described (Rockmill 2009). Spreads were prepared at 24 h for Figures 3a, 5a and S3b and at 15h for Figure S3a. The antibodies used are listed in Table S2. Antibodies were diluted in PBS containing 3% BSA and 10% FBS (Fetal Bovine Serum). Slides were incubated at 4°C overnight (primary antibodies), or at room temperature for 2 hours (secondary antibodies) in a humid chamber. Images of spreads were captured with a Nikon Eclipse 90i fluorescence microscope controlled with MetaMorph software (Molecular Devices) and equipped with a Hammamatsu Orca-AG charge-coupled device (CCD) camera and a PlanApo VC 100x 1.4 NA objective, except for Figure 5a, which were captured with an Olympus IX71 fluorescence microscope equipped with a personal DeltaVision system, a CoolSnap HQ2 (Photometrics) camera, and 100x UPLSAPO 1.4 NA objective. To determine Pch2 and Hop1 intensity on chromosome spreads (Figures 3d, 5b and 5c), a region containing DAPI-stained chromatin was defined and the mean intensity values were measured. The nucleolar region (Figures 3b and 3c) was identified based on the Pch2 signal, and the chromosomal region (Figure 3c) was defined by subtracting the nucleolar region from the total DAPI-stained area. Background values were subtracted using the rolling ball algorithm from Fiji setting the radius to 50 pixels. Images of whole live cells expressing *GFP-PCH2* or *HOP1-GFP* were captured with the Olympus IX71 fluorescence microscope with DeltaVision system described above. For GFP-Pch2, Z-stacks of 7 planes at 0.8-μm intervals were acquired with an exposure time of 800 ms. Single-plane images are shown for individual cells (Figures 4e-4g) and for *PIL1-GBP* fields (Figure 4g). Maximum-intensity projections of three planes are shown for fields in Figures 4e, 4f, and S1, whereas projections of two planes are shown for individual cells in Figure S1. To determine the nuclear/cytoplasm GFP fluorescence ratio (Figures 4h and S1), the ROI manager tool of the Fiji software (Schindelin et al. 2012) was used to define the regions corresponding to the cytoplasm and nucleus (including the nucleolus). Mean fluorescence intensity values were measured from sum-projected images and corrected for background subtraction using the rolling ball algorithm (radius =50 pixels). For Hop1-GFP (Figure 5d), Z-stacks of 14 planes at 0.4-μm intervals were collected with an exposure time of 600 ms. Maximum-intensity projections of selected planes are shown. The nuclear region (including the nucleolus) was defined to measure mean fluorescence intensity from sum-projected images, which were also subjected to background subtraction as described above.

**Figure 4.**
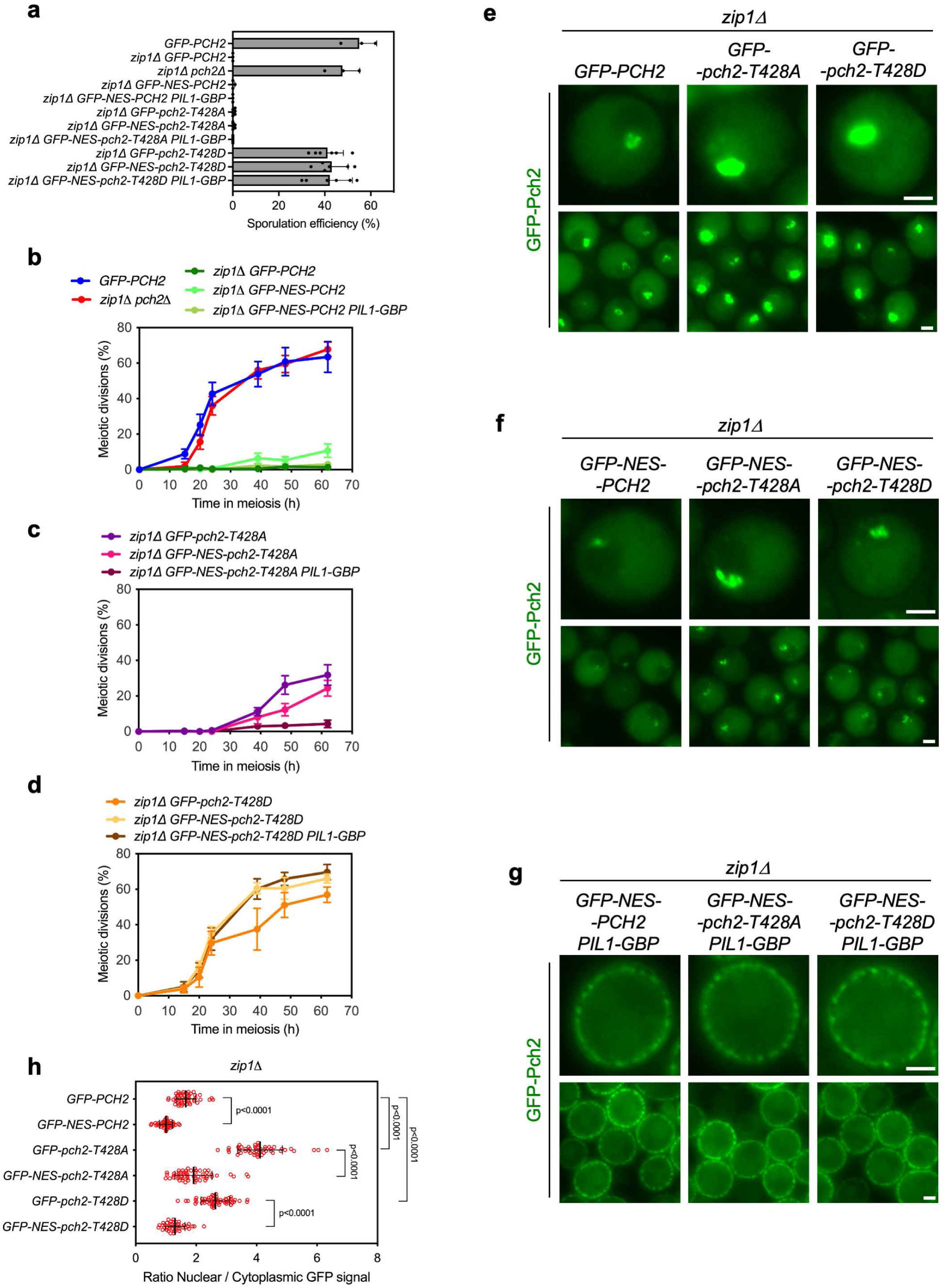
Forced nuclear exit restores checkpoint function for Pch2-T428A, but not Pch2-T428D. **a)** Sporulation efficiency, assessed by microscopic counting of asci, was examined after 3 days on sporulation plates. Error bars, SD; n >= 3. At least 300 cells were counted for each strain. **b-d)** Time-course analysis of meiotic nuclear divisions; the percentage of cells containing two or more nuclei is represented. Error bars: SD; n=3. At least 300 cells were scored for each strain at every time point. **e-g)** Fluorescence microscopy images of the localization of GFP-Pch2 (or its variants) in live meiotic cells of the indicated genotypes. Images were taken 16 h after meiotic induction. Representative individual cells and fields are shown. Scale bar, 2 μm. **h)** Quantification of the ratio of nuclear (including nucleolar) to cytoplasmic GFP fluorescent signal for the experiment shown in (e and f). Error bars, SD. The strains used carry GFP-tagged wild-type *PCH2*, or its mutants derivatives, in hemizygosis. Strains are: DP1624 (*GFP-PCH2*), DP1625 (*zip1Δ GFP-PCH2*), DP1029 (*zip1Δ pch2Δ*), DP1686 (*zip1Δ GFP-NES-PCH2*), DP1796 (*zip1Δ GFP-NES-PCH2 PIL1-GBP*), DP1955 (*zip1Δ GFP-pch2-T428A*), DP1959 (*zip1Δ GFP-NES-pch2-T428A*), DP1964 (*zip1Δ GFP-NES-pch2-T428A PIL1-GBP*), DP1957 (*zip1Δ GFP-pch2-T428D*), DP1961 (*zip1Δ GFP-NES-pch2-T428D*), DP1965 (*zip1Δ GFP-NES-pch2-T428D PIL1-GBP*).

**Figure 5.**
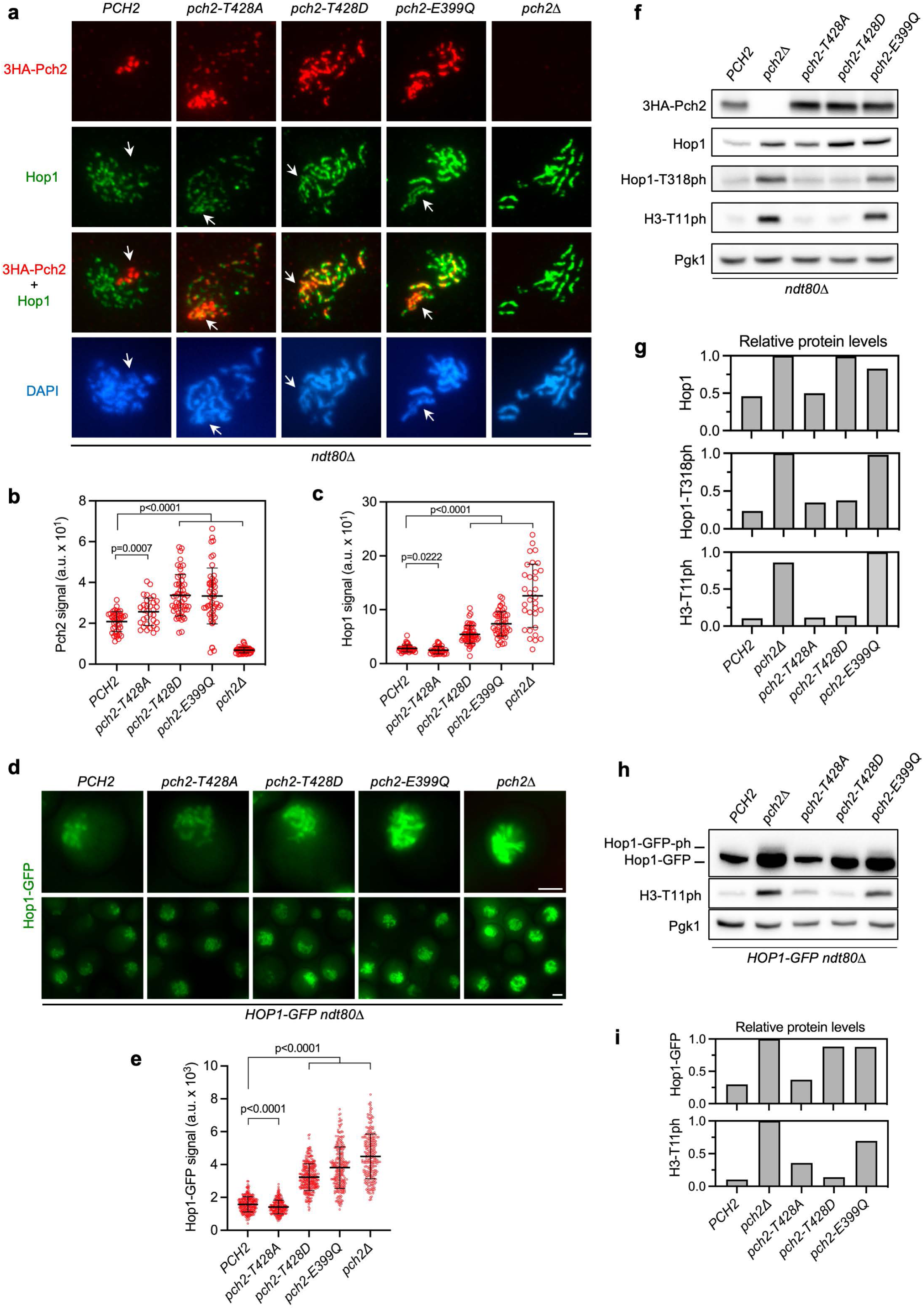
Pch2-T428D dissociates Hop1 localization and phosphorylation. **a)** Immunofluorescence of spread meiotic chromosomes from prophase-I arrested *ndt80*Δ strains, stained with anti-HA (to detect 3HA-Pch2; red) and anti-Hop1 antibodies (green), and with DAPI (blue). Representative nuclei are shown. Spreads were prepared 24 h after meiotic induction. Arrows point to the rDNA region. Scale bar, 2 μm. **b and c)** Quantification of Pch2 and Hop1 signal for the experiment shown in (a). Error bars, SD; a.u., arbitrary units. **d)** Fluorescence microscopy images of the localization of Hop1-GFP in live meiotic cells of the indicated genotypes. Images were taken at 24 h. **e)** Quantification of the Hop1 nuclear signal for the experiment shown in (d). Error bars, SD; a.u., arbitrary units. **f)** Western blot analysis of Pch2 (detected with anti-HA antibodies), Hop1, Hop1-T318ph and H3-T11ph from the same strains analyzed in (a). Pgk1 was used as a loading control. **g)** Quantification of Hop1, Hop1-T318ph and H3T11ph relative levels for the experiment shown in (f). **h)** Western blot analysis of Hop1-GFP (detected with anti-GFP antibodies) and H3-T11ph. The slow-migrating phosphorylated form (Hop1-GFPph) is also marked. Pgk1 was used as a loading control. **i)** Quantification of Hop1-GFP and H3T11ph relative levels for the experiment shown in (h). Strains in (a-c) and (f-g) are: DP1191 (wild type; *3HA-PCH2*), DP1058 (*pch2Δ ndt80Δ*), DP1284 (*3HA-pch2-T428A ndt80Δ*), DP1399 (*3HA-pch2-T428D ndt80Δ*), DP1301 (*3HA-pch2-E399Q ndt80Δ*). Strains in (d-e) and (h-i) are: DP2375 (wild type; *3HA-PCH2 ndt80Δ HOP1-GFP/HOP1*), DP2376 (*3HA-pch2-T428A ndt80Δ HOP1-GFP/HOP1),* DP2377 (*3HA-pch2-T428D ndt80Δ HOP1-GFP/HOP1*), DP2378 (*3HA-pch2-E399Q ndt80Δ HOP1-GFP/HOP1*), DP2374 (*pch2Δ ndt80Δ HOP1-GFP/HOP1)*.

### Sporulation efficiency and spore viability

Sporulation efficiency was quantitated by microscopic examination of asci formation after 3 days on sporulation plates. Both mature and immature asci were scored. At least 300 cells were counted for every strain. Spore viability was assessed by tetrad dissection. At least 144 spores were scored for every strain.

### Software and statistics

The predicted structure of yeast Pch2 (UniProt P38126) was obtained from the AlphaFold3 server (Abramson et al. 2024). The UCSF Chimera software was used to visualize and annotate the protein structure model (Meng et al. 2006). Statistical analyses were performed using GraphPad Prism version 11.0. Pairwise comparisons were performed using unpaired two-tailed tests. Normality was assessed with the Shapiro-Wilk test. For normally distributed data, homogeneity of variance was evaluated using an F-test for equality of variances. Student’s *t*-test was used when variances were equal, and Welch’s *t*-test when variances were unequal. When data did not follow a normal distribution, comparisons were performed using the Mann-Whitney U test. The type of error bars and the number of biological replicates are indicated in the corresponding figure legends, when applicable. Detailed results of statistical analyses, including all pairwise comparisons and corresponding *P* values for each relevant figure panel, are provided in Supplementary Data 1. Raw data underlying the graphs are available in Supplementary Data 2.

## RESULTS

### Pch2 contains a conserved Mec1/Tel1 consensus phosphorylation site

To investigate the possible regulation of Pch2 function or localization by phosphorylation, we searched the budding yeast Pch2 protein sequence for potential phosphorylation motifs and identified a consensus site (S/T-Q) for the Mec1/Tel1 checkpoint kinases (Mallory and Petes 2000) at position 428 (Figure 1a). This motif, located in the C-terminal AAA+ domain, is conserved in Pch2 orthologs from *C. elegans* (PCH-2) and humans (TRIP13) (Figure 1b), suggesting its functional importance. Moreover, electrophoretic analysis of the Pch2 protein in Phos-tag gels occasionally revealed two slow-migrating forms of Pch2 that are suggestive of phosphorylation. We observed that the upper band was greatly reduced in *mec1*Δ and *rad24*Δ mutants (Figure 1c). Thus, to explore the biological relevance of this TQ motif for Pch2 meiotic checkpoint function, we mutated the Thr428 residue to alanine and aspartic acid, to generate the putative phosphodeficient (*pch2-T428A*) and phosphomimetic (*pch2-T428D*) mutants, respectively (Figure 1a).

We found that the upper band was also less evident in *pch2-T428A* and in *rad24*Δ *tel1*Δ mutants, similar to the pattern detected in *mec1*Δ and *rad24*Δ, but it remained present in the *tel1*Δ single mutant (Figure 1d). We note that because Rad24 acts on the Mec1 signaling branch, we used *rad24*Δ *tel1*Δ to abolish both the Mec1 and Tel1 pathways. This approach was followed because the *mec1*Δ *tel1*Δ double mutant exhibits poor viability even in a *sml1*Δ background, which precludes an accurate analysis. These observations suggest that the T428 residue of Pch2 could be a target for Mec1-dependent phosphorylation. We also observed that the slower mobility forms were completely abolished in the *pch2-K320A* mutant, which carries an amino acid change in the ATP binding site (Walker A, Figure 1a). Since the Pch2-K320A protein is unable to form hexamers, and its chromatin association is impaired (Herruzo et al. 2016), it is possible that the putative Pch2 post-translational modification of Pch2 occurs in the chromosomal and/or nucleolar population. Nevertheless, we point out that this is currently the only evidence for Pch2-T428 phosphorylation. For unknown technical reasons, we have not been able to systematically replicate this mobility change. Despite that, because this TQ motif is evolutionarily conserved, we proceeded with its functional characterization independently of its phosphorylation status.

### The meiotic checkpoint response is altered to different extents in *pch2-T428A* and *pch2-T428D* mutants

We first examined the relevance of Pch2-Thr428 in a *zip1*Δ mutant background where Pch2 is critically required for the checkpoint response (San-Segundo and Roeder 1999; Herruzo et al. 2016). The strong sporulation defect of *zip1*Δ was either partially or completely relieved in the *zip1*Δ *pch2-T428A* or *zip1*Δ *pch2-T428D* double mutants, respectively, originating spores with decreased viability (≅45-50%), as expected when the checkpoint arrest is bypassed (Figure 2a). Likewise, time-course analysis of meiotic nuclear divisions revealed that the pronounced meiotic progression delay displayed by *zip1*Δ was either partially or completely abolished in the *zip1*Δ *pch2-T428A* or *zip1*Δ *pch2-T428D* double mutants, respectively (Figure 2b). We also evaluated meiotic recombination checkpoint activity during these meiotic time courses by monitoring molecular markers of checkpoint signaling, including phosphorylation of Hop1-T318 (a Mec1 target), H3-T11 (a Mek1 target), and Mek1 itself (Carballo et al. 2008; Chuang et al. 2012; Kniewel et al. 2017). As previously described, *zip1*Δ exhibited high and persistent levels of phosphorylation of these checkpoint markers; only at late time points, checkpoint activity mildly declined coincident with resumption of meiotic progression in a subset of *zip1*Δ cells (Figure 2b, 2c). Consistent with the kinetics of nuclear divisions (Figure 2b), checkpoint signaling was only slightly impaired in the *zip1*Δ *pch2-T428A* double mutant, with Hop1-T318, Mek1 and H3-T11 phosphorylation levels only marginally attenuated and disappearing only a the very late time point (Figure 2c). In contrast, the *zip1*Δ *pch2-T428D* mutant exhibited a pronounced decline in checkpoint activity, with the dynamics of checkpoint markers similar to the wild type (Figure 2c), consistent with the full suppression of the *zip1*Δ meiotic delay (Figure 2b). Since Pch2 is produced during meiotic prophase I, the dynamics of Pch2 levels correlated with the extent of the checkpoint-dependent prophase I arrest; that is, high levels of Pch2 largely persisted in the *zip1*Δ mutant, they were slightly reduced at late time points in *zip1*Δ *pch2-T428A*, and they substantially dropped in the *zip1*Δ *pch2-T428D* mutant as the meiotic block was released (Figure 2b, 2c). These observations indicate that checkpoint function is only mildly affected in the *zip1*Δ *pch2-T428A* mutant, but it is severely compromised in *zip1*Δ *pch2-T428D*.

### Hop1 localization to unsynapsed *zip1*Δ axes is impaired in *pch2-T428D*

In nuclear spreads of the *zip1*Δ mutant, Hop1 continuously decorates chromosome axes; this linear localization requires Pch2 function. In fact, in contrast to the *pch2*Δ single mutant, which shows increased levels of Hop1 on the SC, in the *zip1*Δ *pch2*Δ or *zip1*Δ *pch2-E399Q* double mutants, Hop1 axial localization is reduced and discontinuous (Figure 3a, 3d) (Herruzo et al. 2016). Therefore, to determine whether T428 is important for Hop1 localization on unsynapsed *zip1*Δ chromosomes, we examined Hop1 and Pch2 distribution on spreads of the *zip1*Δ *pch2-T428A* and *zip1*Δ *pch2-T428D* double mutants. We performed this analysis in prophase I-arrested *ndt80*Δ strains to avoid interference with the different rates of cell cycle progression among the various mutants tested (Figure 2b). Like wild-type Pch2, both Pch2-T428A and Pch2-T428D versions showed an exclusive nucleolar localization in *zip1*Δ and the intensity of the Pch2 signal did not significantly vary among the different versions (Figure 3a, 3b). However, whereas Hop1 was absent from the rDNA region covered by wild-type Pch2; it was weakly detectable in the nucleolar area of *zip1*Δ *pch2-T428A* (Figure 3a). Furthermore, Hop1 clearly decorated the rDNA axes of *zip1*Δ *pch2-T428D*, colocalizing with Pch2-T428D, as occurs in *zip1*Δ *pch2-E399Q*. These observations suggest that the checkpoint-independent activity of Pch2 to ensure Hop1 exclusion from the rDNA array is impaired in T428 mutants to different extents, as revealed by quantifying the ratio between nucleolar and chromosomal Hop1 signals (Figure 3c).

With regard to chromosomal Hop1, the *zip1*Δ *pch2-T428A* double mutant exhibited a continuous linear pattern comparable to that of the *zip1*Δ single mutant (Figure 3a). In contrast, *zip1*Δ *pch2-T428D* showed a disrupted Hop1 distribution, displaying reduced linearity and quantity, a pattern similar to that of *zip1*Δ strains lacking Pch2 or expressing a catalytically inactive Pch2 protein (Figure 3a, 3d). This observation is consistent with the strong checkpoint defect of *zip1*Δ *pch2-T428D*.

We have previously shown that the crucial checkpoint role of Pch2 is to sustain Hop1 phosphorylation at T318 (Herruzo et al. 2016). Since Pch2 also controls global levels of Hop1, we measured the ratio between Hop1-T318ph and total Hop1 in cell extracts from *zip1*Δ *ndt80*Δ-arrested cells to assess Pch2 checkpoint function (Herruzo et al. 2019). Mek1-dependent H3-T11ph, which depends on prior phosphorylation of Hop1 at T318, was also analyzed (Carballo et al. 2008; Chuang et al. 2012; Kniewel et al. 2017) (Figure 3e). The *zip1*Δ *pch2-T428A* mutant showed only slightly decreased Hop1-T318ph/Hop1 ratio and quite high levels of Mek1 activity indicating that the checkpoint function of Pch2-T428A was only marginally affected (Figure 3e, 3f). On the other hand, similar to *zip1*Δ *pch2*Δ and *zip1*Δ *pch2-E399Q*, checkpoint activity was almost completely abolished in *zip1*Δ *pch2-T428D*, as manifested by the strongly reduced Hop1-T318ph/Hop1 ratio and diminished Mek1 signaling (Figure 3e, 3f). In sum, these findings indicate that the presence of a negative charge at position 428 of Pch2 severely impairs its ability to promote high levels of Hop1-T318ph to sustain the *zip1*Δ-induced checkpoint response.

### Pch2-Thr428 is required for proper subcellular distribution

Besides the nucleolus and synapsed chromosomes, Pch2 is also localized in the cytoplasm of meiotic prophase I cells; in fact, a balanced subcellular distribution of Pch2 between nuclear and cytoplasmic compartments is critical for precise checkpoint activity. To explore whether the *pch2-T428A* and *pch2-T428D* mutations affect Pch2 subcellular localization, we generated strains harboring GFP-tagged versions expressed from the *HOP1* promoter as previously described (Herruzo et al. 2019). For simplicity, these constructs will be referred to as *GFP-PCH2* (or mutant derivatives) throughout this article. To assess *zip1*Δ-induced checkpoint function we examined sporulation efficiency as well as kinetics of meiotic nuclear divisions. Like *zip1*Δ *GFP-PCH2*, the *zip1*Δ *GFP-pch2-T428A* double mutant displayed a strong sporulation block (Figure 4a), and showed inefficient and delayed meiotic divisions (Figure 4b, 4c), indicating that the checkpoint remained active to some extent. In contrast, *zip1*Δ *GFP-pch2-T428D* completely suppressed the sporulation arrest (Figure 4a), and exhibited a kinetics of meiotic progression comparable to that of *zip1*Δ *pch2*Δ (Figure 4b, 4d), confirming that checkpoint function is seriously impaired when Pch2-Thr428 is changed to Asp.

We next examined localization of the GFP-tagged mutant versions in live meiotic cells. In the *zip1*Δ mutant, the wild-type GFP-Pch2 protein is exclusively present in the cytoplasm and nucleolus, and no signal is detected in the rest of the nucleus, as described (Herruzo et al. 2019; Herruzo et al. 2021; Herruzo et al. 2023) (Figure 4e). In contrast, the nucleolar signal was much stronger in the *zip1*Δ *GFP-pch2-T428A* and *zip1*Δ *GFP-pch2-T428D* mutants (Figure 4e). In the *zip1*Δ *GFP-pch2-T428A,* the rest of the nucleus also exhibited a diffuse signal or even occasional foci (Figure 4e). Indeed, the nuclear/cytoplasmic fluorescence ratio was higher in *zip1*Δ *GFP-pch2-T428A* and *zip1*Δ *GFP-pch2-T428D* compared to *zip1*Δ *GFP-PCH2* strains (Figure 4h). Thus, the balanced subcellular distribution of Pch2 is disrupted when T428 is mutated.

Since the equilibrium between the nuclear and cytoplasmic Pch2 pools dictates a proper checkpoint response and the increased accumulation of Pch2 in the nucleus impairs it (Herruzo et al. 2021; Herruzo et al. 2023), the biased distribution of Pch2-T428A and Pch2-T428D towards the nucleus may solely explain the defective checkpoint in *zip1*Δ *GFP-pch2-T428A* and *zip1*Δ *GFP-pch2-T428D* mutants (Figure 4a-4d). Alternative, or in addition, the mutations could compromise other aspects of Pch2 biology, such as the enzymatic activity of the proteins or the interaction with other partners, thus explaining the defective checkpoint function. To determine whether the checkpoint phenotype of the T428 mutants is exclusively due to the aberrant subcellular distribution of Pch2 we reverted the nuclear accumulation. We forced the localization outside the nucleus in two ways: 1) fusion to a strong nuclear export signal (NES) (Figure 4f), and 2) tethering to the plasma membrane via the binding of GFP to the eisosome component Pil1 tagged with the GFP-binding protein (GBP) (Figure 4g), as previously described (Herruzo et al. 2021).

The addition of a strong NES to either Pch2-T428A or Pch2-T428D led to a dramatic decrease in the nuclear/cytoplasmic ratio, with a weaker nucleolar signal remaining (Figure 4f, 4h). In the case of *zip1*Δ *NES-pch2-T428A* the checkpoint function was largely restored, as manifested by the lack of sporulation (Figure 4a) and the more pronounced delay in meiotic progression (Figure 4c). However, the checkpoint was still defective in *zip1*Δ *NES-pch2-T428D*, which displayed high sporulation efficiency (Figure 4a) and kinetics of meiotic progression similar to that of *zip1*Δ *pch2*Δ (Figure 4b, 4d). Likewise, Pil1-GBP-mediated tethering of Pch2-T428A to the cell periphery restored checkpoint function (Figure 4a-4c), but tethering Pch2-T428D did not (Figure 4a, 4b, 4d). These results imply that while the modest checkpoint defect conferred by the substitution of Thr428 to Ala primarily results from the abnormal accumulation of the protein in the nucleus; the function of the Pch2-T428D version is altered beyond the anomalous localization.

### Mutation of Thr428 affects Pch2 and Hop1 chromosomal localization in synapsis proficient strains

The results described above indicate that Pch2-T428 is involved in the checkpoint response triggered by *zip1Δ*. Next, we analyzed the effect of T428 mutations during unperturbed meiosis by examining Pch2 and Hop1 localization on prophase I chromosome spreads from synapsis proficient *ZIP1* strains. In the wild type, Pch2 protein massively accumulates in the unsynapsed rDNA array on chromosome XII, where it prevents Hop1 localization and meiotic DSB formation (Figure 5a) (San-Segundo and Roeder 1999; Vader et al. 2011; Herruzo et al. 2019; Villar-Fernández et al. 2020). Pch2 also localizes to foci on synapsed chromosomes, although these chromosomal foci are often barely detectable in the BR1919 strain background (San-Segundo and Roeder 1999; Herruzo et al. 2021). We found that the nucleolar localization of Pch2-T428A was also quite robust, but the chromosomal foci were more conspicuous compared to the wild type. In contrast, Pch2-T428D displayed an extensive distribution throughout the chromatin, showing a linear pattern comparable to that of the catalytically inactive Pch2-E399Q protein (Figure 5a, 5b). Analysis of the nuclear to cytoplasmic ratio of GFP-tagged Pch2 versions in *ZIP1* cells also revealed an increased accumulation of Pch2-T428A in the nucleus, particularly concentrated in the nucleolus, and confirmed the substantial chromosomal localization of Pch2-T428D (Figure S1).

In turn, the localization of Hop1 in the *pch2-T428A* mutant was similar to the wild type, displaying a limited distribution on synapsed chromosomes. Conversely, despite the abundance of chromosomal Pch2-T428D, Hop1 accumulated to high levels in the *pch2-T428D* mutant, although it did not reach the levels observed in *pch2-E399Q* or *pch2Δ* (Figure 5a, 5c). To reinforce these observations, we also examined the localization of a C-terminal GFP-tagged version of Hop1. We previously confirmed that, in an otherwise wild-type background, Hop1-GFP supports high spore viability, which is indicative of the protein’s functionality in this context, thus validating its use in *ZIP1* strains (Figure S2a). We note, however, that Hop1-GFP did not support the checkpoint arrest of *zip1Δ* (Figure S2b). Analysis of Hop1-GFP localization in live meiotic cells confirmed that its distribution was not substantially altered in *pch2-T428A*, showing only slightly reduced intensity. However, it displayed a progressively increased signal in the *pch2-T428D*, *pch2-E399Q* and *pch2Δ* mutants, respectively (Figure 5d, 5e). Together, these observations indicate that while both T428 mutations increase Pch2 localization on chromosome axes, their functional impacts differ: the ability to remove Hop1 from the synapsed chromosomes remains largely intact in Pch2-T428A, whereas it is significantly impaired in Pch2-T428D.

Pch2 chromosomal recruitment is known to depend on Zip1, suggesting an interaction between both proteins in the SC, as has been described in plants (Yang et al. 2022). This Pch2-Zip1 association is clearly manifested by their extensive colocalization within polycomplexes, which are large extrachromosomal aggregates of SC components (Sym et al. 1993; Sym and Roeder 1995; San-Segundo and Roeder 1999; Borner et al. 2008). Therefore, to determine whether mutation of T428 affects the Pch2-Zip1 interaction, we examined the recruitment of Pch2 to polycomplexes in strains overexpressing *ZIP1* to induce these structures. Like the wild-type protein, both Pch2-T428A and Pch2-T428D colocalized with Zip1 in these organized SC aggregates (Figure S3a), suggesting that the interaction with Zip1 persists in the T428 mutants. Furthermore, analysis of Zip1 and Pch2 localization on synapsed chromosomes showed that the Zip1 linear pattern was not altered in T428 mutants. However, a substantial overlap between Zip1 and Pch2-T428D was observed, consistent with an increased recruitment and/or maintenance of Pch2-T428D on synapsed chromosomes (Figure S3b).

### The *pch2-T428D* mutation uncouples Hop1 accumulation from its phosphorylation

Pch2 activity influences not only the chromosomal distribution of Hop1, but also the global levels of Hop1 protein and its phosphorylation, specifically at the T318 residue. In contrast to *zip1Δ*, in synapsis proficient *ZIP1* strains lacking Pch2 activity, such as *pch2Δ* or *pch2-E399Q*, both Hop1 and Hop1-T318ph levels are increased (Herruzo et al. 2016) (Figure 5f). We therefore performed western blot analysis to determine whether the T428 mutations of Pch2 affect Hop1 and Hop1-T318ph levels in prophase I arrested *ndt80Δ* strains. We also examined H3-T11ph as a readout for Mek1 activity, which depends on the prior phosphorylation of Hop1-T318. The *pch2-T428A* mutant did not show substantial differences from the wild type (Figure 5f, 5g). In contrast, the *pch2-T428D* mutant exhibited higher global Hop1 protein levels. However, unlike the *pch2-E399Q* or *pch2Δ* mutants, this increase was not accompanied by elevated levels of Hop1-T318ph or H3-T11ph (Figure 5f, 5g). Likewise, analysis of *HOP1-GFP* strains also revealed that the increased amount of Hop1-GFP protein in *pch2-T428D* was not paralleled by increased phosphorylation, detected as a slow-migrating form, resulting in low H3-T11ph levels (Figure 5h, 5i). Thus, while *pch2-T428D* phenocopies loss-of-function mutants in a *zip1Δ* background, it does not do so in *ZIP1* strains. This suggests that the functional consequences on Hop1 dynamics are not solely explained by a simple lack of Pch2 activity.

Taken together, these results reveal that the T428D substitution in Pch2 specifically interferes with the ability of Hop1 to undergo phosphorylation, despite its increased accumulation on chromosome axes. By genetically separating these two events, our findings indicate that the presence of negative charge at Pch2-T428, which may mimic phosphorylation, is sufficient to modify Hop1-Mek1 signaling dynamics.

### The *pch2-T428D* mutation compromises meiotic chromosome segregation when DSBs are limiting

Given that Hop1-T318 phosphorylation and subsequent Mek1 activation are required to ensure accurate meiotic recombination, we investigated whether the altered signaling observed in *pch2-T428D* had functional consequences for the completion of meiosis. Sporulation efficiency and meiotic progression were normal in *pch2-T428A* and *pch2-T428D* single mutants (Figure 6a, 6b), and spore viability was minimally affected (Figure 6c). This is consistent with the fact that Pch2 function is largely dispensable during unperturbed meiosis. In fact, spore viability, an indirect measure of meiotic chromosome segregation fidelity, is largely unaffected in the *pch2*Δ or *pch2-E399Q* single mutants (San-Segundo and Roeder 1999; Borner et al. 2008; Chen et al. 2014). However, it becomes compromised in sensitized situations where DSB levels are reduced, such as when combined with a *spo11-3HA* hypomorphic allele (Zanders and Alani 2009) (Figure 6c). We found that spore viability was fairly normal in *spo11-3HA pch2-T428A*, but it was mildly reduced in *spo11-3HA pch2-T428D,* although to a lesser extent than that observed in *spo11-3HA pch2*Δ or in *spo11-3HA pch2-E399Q* (Figure 6c). The *spo11-3HA pch2-T428D* mutant displayed a moderate bias toward 4, 2, and 0 viable spore tetrads, which is indicative of meiosis I nondisjunction events (Figure 6d). These observations suggest that the altered coordination of Hop1 loading and phosphorylation caused by the presence of a negative charge at Pch2-T428 becomes detrimental for meiotic success when DSB formation is suboptimal.

**Figure 6.**
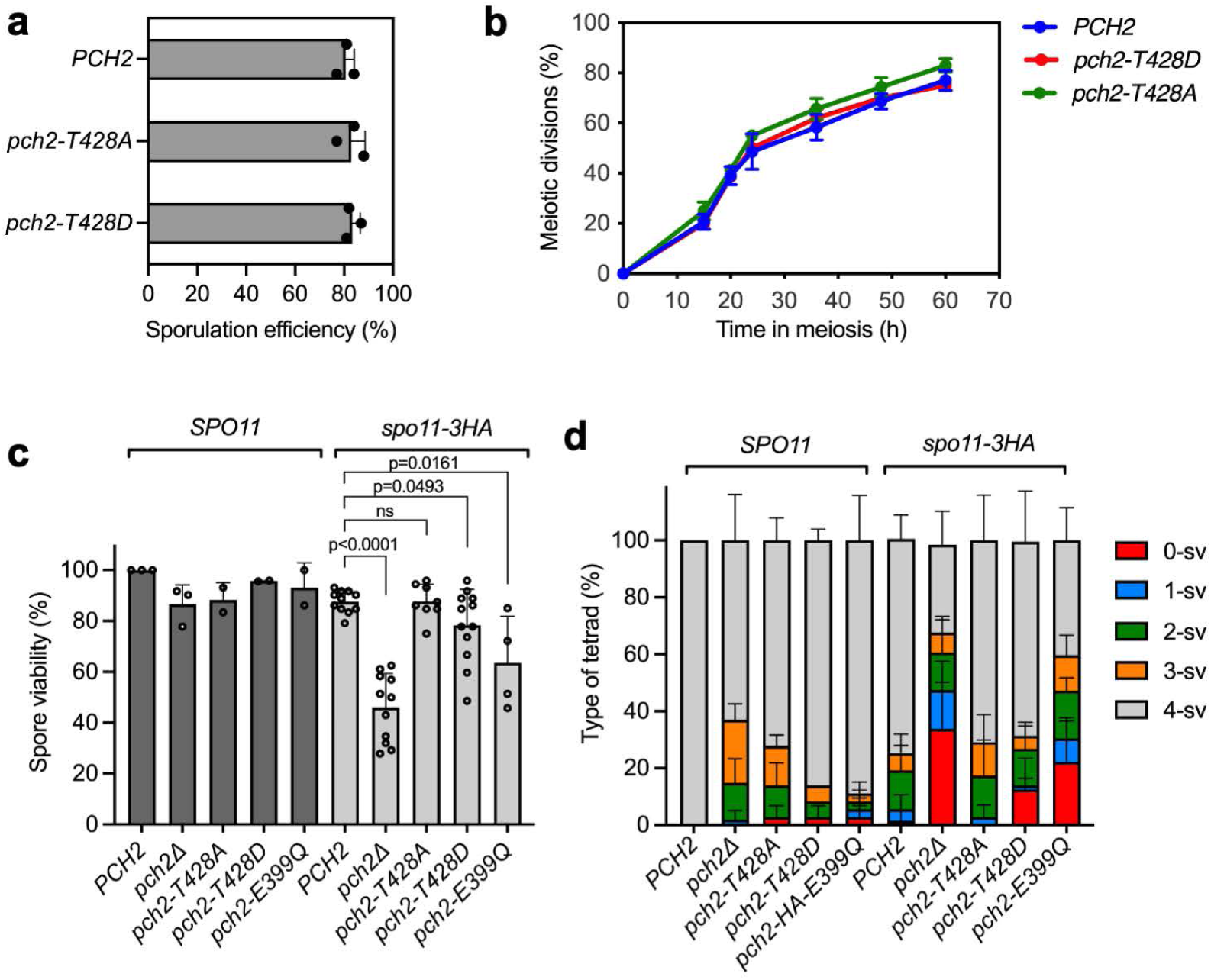
The *pch2-T428D* mutation reduces spore viability when DSB formation is limiting. **a)** Sporulation efficiency, assessed by microscopic counting of asci, was examined after 3 days on sporulation plates. Error bars, SD; n = 3. At least 300 cells were counted for each strain. **b)** Time-course analysis of meiotic nuclear divisions; the percentage of cells containing two or more nuclei is represented. Error bars: SD; n=3. At least 300 cells were scored for each strain at every time point. Strains in (a-b) are: DP1151 (wild type; *3HA-PCH2*), DP1256 (*3HA-pch2-T428A*), DP1332 (*3HA-pch2-T428D*). **c)** Spore viability assessed by tetrad dissection. **d)** percentage of tetrads containing 4-, 3-, 2-, 1-, and 0-viable spores. At least 36 tetrads were dissected for each strain. Error bars, SD. Strains in (c) and (d) are: DP1151 (wild type; *3HA-PCH2*), DP1023 (*pch2Δ*), DP1256 (*3HA-pch2-T428A*), DP1332 (*3HA-pch2-T428D*), DP1287 (*3HA-pch2-E399Q*), DP1948 (*spo11-3HA 3HA-PCH2*), DP1787 (*spo11-3HA pch2Δ),* DP1949 (*spo11-3HA 3HA-pch2-T428A*), DP1950 (*spo11-3HA 3HA-pch2-T428D*), and DP2074 (*spo11-3HA 3HA-pch2-E399Q*).

## DISCUSSION

In this study, we investigated the functional impact of threonine 428 (T428) in the Pch2 protein, a conserved AAA+ ATPase critical for meiotic chromosome dynamics. T428 lies within a threonine-glutamine (TQ) motif conserved among Pch2 orthologs and represents a potential target for Mec1/Tel1 checkpoint kinases. We initially hypothesized that phosphorylation of Pch2-T428 could regulate its role in the meiotic recombination checkpoint. This idea was also supported by the known physical interaction between Pch2 and the region of Xrs2 containing BRCT repeats, which are modules that typically recognize phosphorylated motifs (Yu et al. 2003; Becker et al. 2006; Ho and Burgess 2011). In addition, initial Phos-tag experiments suggested the presence of slower-migrating Pch2 species that were reduced in *pch2-T428A* and *mec1Δ* mutants, raising the possibility of modification at this site; however, this pattern could not be reproducibly detected. Although direct evidence for T428 phosphorylation remains inconclusive, our mutational analysis clearly indicates that this residue is functionally important for Pch2-dependent regulation of meiosis, both in checkpoint-activated (*zip1Δ*) and unperturbed (*ZIP1*) contexts, as summarized in Figure 7.

**Figure 7.**
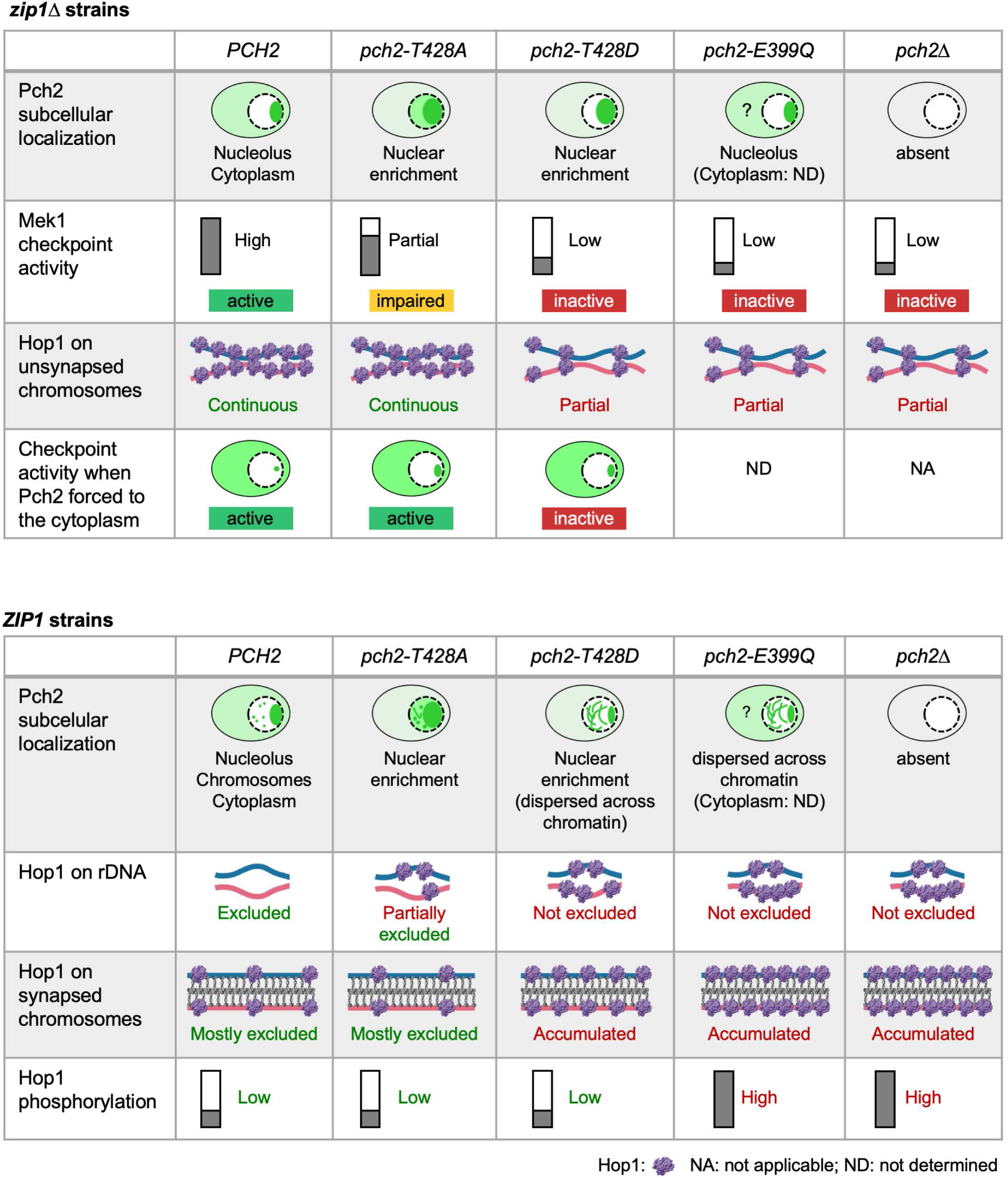
Schematic summary of Pch2 functional and localization analyses in *zip1Δ* and *ZIP1* strains. Graphical overview of the cytological and molecular phenotypes associated with distinct *PCH2* alleles in different synapsis contexts, including protein localization, chromosome organization and checkpoint signaling. See text for details.

To evaluate how the T428 residue influences checkpoint control, we monitored the ability of the *pch2-T428A* and *pch2-T428D* mutants to maintain the *zip1Δ*-induced meiotic delay and assessed Mec1-dependent Hop1-T318ph and Mek1-dependent H3-T11ph, two established readouts of Pch2 checkpoint activity (Cavero et al. 2016). The checkpoint appears only mildly compromised in *zip1Δ pch2-T428A*; in contrast, *zip1Δ pch2-T428D* exhibits a pronounced checkpoint defect that closely resembles the phenotype of *zip1Δ pch2Δ* (Herruzo et al. 2016). These findings suggest that T428 contributes to Pch2’s checkpoint role and that, irrespective of whether this residue undergoes phosphorylation in vivo, introducing a negative charge at this position interferes with proper Pch2 function.

Although the nucleolar, rDNA-associated population of Pch2 is not directly required for checkpoint control (Herruzo et al. 2019), the differential impact of the T428A and T428D substitutions is clearly evident in their effects on Hop1 exclusion from the rDNA. In *zip1Δ pch2-T428A* cells, Hop1 shows only a weak nucleolar signal, suggesting that Pch2-T428A retains substantial ability to prevent Hop1 accumulation in the rDNA array. In contrast, Hop1 heavily decorates the rDNA region in *zip1Δ pch2-T428D* mutants, revealing a pronounced defect in this aspect of Pch2 function. Because Hop1 exclusion from the rDNA likely relies on Pch2-driven remodeling events that influence Hop1 conformation, localization, or turnover, the strong nucleolar accumulation observed in *pch2-T428D* chromosome spreads reflects a major impairment in the capacity of the Pch2-T428D protein to regulate Hop1 dynamics.

Live-cell analysis of GFP-tagged Pch2 in *zip1Δ* strains reveals that substitution of T428 disrupts the balanced nucleocytoplasmic distribution required for a proper checkpoint response (Raina and Vader 2020; Herruzo et al. 2021). Notably, Pch2 nuclear accumulation is known to impair the checkpoint, whereas trapping Pch2 in an extranuclear compartment enhances it (Herruzo et al. 2021). Both Pch2-T428A and Pch2-T428D accumulate excessively and abnormally within the nucleus, suggesting that mutation of this residue interferes with Pch2 nuclear export. This mislocalization appears to account for the mild checkpoint defect of *zip1Δ pch2-T428A*, as forcing this mutant protein to remain outside the nucleus fully restores checkpoint activity. In contrast, Pch2-T428D behaves differently: although it also exhibits an abnormal nuclear enrichment, redirecting it to an extranuclear compartment fails to rescue its checkpoint defect. This result uncouples the localization defect from the functional defect, arguing that a negative charge at T428 disrupts an additional, localization-independent activity of Pch2. This is consistent with the more severe phenotypes of *zip1Δ pch2-T428D*, which mirror those of *zip1Δ pch2Δ*. Because Pch2 undergoes Crm1-dependent nucleocytoplasmic trafficking via an internal nuclear export sequence (NES), a process required for a proper checkpoint response (Herruzo et al. 2023), the increased nuclear retention of both T428 mutants suggests a defect in this trafficking event, possibly through altered NES accessibility or recognition.

Analysis of T428 mutant phenotypes in synapsis-proficient strains also revealed critical functional distinctions. While both mutant proteins exhibit increased nuclear accumulation, Pch2-T428D shows a pronounced enrichment along synapsed chromosomes, where it extensively colocalizes with Zip1 and Hop1. This contrasts with the more limited chromosomal association of wild-type Pch2 and Pch2-T428A, indicating that a negative charge at T428 perturbs not only nucleocytoplasmic trafficking, but also the dynamic binding and/or release of Pch2 from the SC, which shapes the regional organization of Hop1 and Zip1 along the SC (Borner et al. 2008; Subramanian et al. 2019). Such excessive recruitment or retention of Pch2-T428D may underlie the mildly reduced spore viability observed when this mutant is combined with a hypomorphic *spo11-3HA* allele. Under low-DSB conditions, where recombination intermediates must be precisely managed, inappropriate Pch2 retention along the SC may disrupt Hop1 remodeling, axis organization, and recombination homeostasis.

T428 lies within a conserved region of the AAA+ ATPase module, spatially close to elements critical for ATP hydrolysis (Walker B) and for conformational transitions of the hexameric ring required for substrate remodeling (Hanson and Whiteheart 2005; Wendler et al. 2012; Puchades et al. 2020). AlphaFold-based structural predictions place T428 on the surface of the AAA+ domain where it could influence nucleotide-dependent conformational changes or interactions with regulatory partners (Figure S4). Substituting T428 with a negatively charged residue may subtly alter local structure or electrostatic properties affecting ATP-driven remodeling efficiency or protein-protein interactions. This likely explains why Pch2-T428D exhibits stronger defects than Pch2-T428A. Indeed, several phenotypes of *pch2-T428D,* including the defective checkpoint in *zip1Δ* and the increased Hop1 chromosomal association in *ZIP1* strains, could reflect diminished catalytic efficiency of the Pch2-T428D protein, though direct biochemical measurement of ATPase activity remain necessary to confirm this. However, *pch2-T428D* does not fully mimic the ATPase-dead *pch2-E399Q* or *pch2Δ* mutants in synapsis-proficient contexts. In *pch2Δ* and *pch2-E399Q*, the increase in total Hop1 levels is accompanied by increased Hop1-T318ph and subsequent hyper-active Mek1 signaling (H3-T11ph). In *pch2-T428D*, however, these effects are uncoupled: augmented Hop1 levels do not translate into increased Mek1 activation. This suggests that *pch2-T428D* specifically impairs Pch2’s ability to generate a phosphorylation-competent Hop1-Red1 configuration on chromosome axes, perhaps by failing to correctly present Hop1 to the Mec1 kinase.

These observations collectively suggest that Pch2 contributes to meiotic robustness through separable activities that coordinate Hop1 accumulation, its Red1-dependent phosphorylation, and chromosome-axis organization. In *pch2Δ* or *pch2-E399Q*, the loss of Pch2-mediated restraint leads to hyperactivation of the Hop1-Mek1 signaling pathway that likely disrupts crossover homeostasis and makes the cells particularly sensitive to low-DSB conditions. In contrast, *pch2-T428D* fails to boost Mek1 signaling despite high total Hop1 levels, suggesting that this mutation specifically impairs Pch2’s ability to promote the transition of Hop1 into a phosphorylation-competent state. The milder synthetic spore viability defect of *pch2-T428D spo11-3HA* compared to *pch2Δ spo11-3HA* (or *pch2-E399Q spo11-3HA*) suggests that excessive, misregulated Hop1-Mek1 signaling (*pch2Δ*) is more detrimental to meiotic success than the insufficient signaling observed in *pch2-T428D* under conditions of DSB limitation.

### Concluding Remarks

In summary, our study identifies T428 as a critical residue for the diverse meiotic functions of the AAA+ ATPase Pch2. While the precise biochemical nature of T428 regulation remains to be fully elucidated, our findings demonstrate that this site is essential for orchestrating Pch2’s subcellular distribution and its catalytic impact on chromosomal substrates. The distinct phenotypes of the *T428A* and *T428D* mutants allowed us to uncouple the requirements for nuclear export from the ability of Pch2 to remodel Hop1 into a signaling-competent state. Specifically, our results in *zip1Δ* strains show that T428 is necessary for a functional checkpoint response, while the analysis of synapsis-proficient strains reveals that this residue is also important for the proper coordination of Hop1 levels and Mek1 activation along the SC. By characterizing this regulatory node, we provide further insight into the complex mechanisms that ensure the fidelity of chromosome segregation and the successful completion of the meiotic program.

## Acknowledgements

We are grateful to Andrés Clemente, Raimundo Freire, Jaime Correa, Jesús Carballo, Neil Hunter, and members of the PSS lab for helpful discussions and for reagents. We also acknowledge Jesús Pinto for advice in microscopy image analysis.

## Author contributions

EH: conceptualization, investigation, formal analysis, visualization, writing original draft.

ST: formal analysis.

BS: conceptualization, supervision, project administration.

PSS: conceptualization, investigation, supervision, funding acquisition, visualization, writing original draft, review and editing.

All authors revised, commented, and approved the manuscript.

## Data availability

All relevant data are within the manuscript and the Supplementary information files. Yeast strains generated in this study, as well as any other relevant information are available from the corresponding author upon reasonable request.

